# Revealing the high propensity of RNAs to non-specifically bind drug-like small molecules

**DOI:** 10.1101/2020.05.02.074336

**Authors:** Megan L. Kelly, Chia-Chieh Chu, Honglue Shi, Laura R. Ganser, Hal P. Bogerd, Kelly Huynh, Yuze Hou, Bryan R. Cullen, Hashim M. Al-Hashimi

## Abstract

Identifying small molecules that selectively bind a single RNA target while discriminating against all other cellular RNAs is an important challenge in RNA-targeted drug discovery. Much effort has been directed toward identifying drug-like small molecules that minimize electrostatic and stacking interactions that lead to non-specific binding of aminoglycosides and intercalators to a variety of RNAs. Many such compounds have been reported to bind RNAs and inhibit their cellular activities, however the ability of such compounds to discriminate against RNA stem-loops commonly found in the transcriptome has not been thoroughly assessed in all cases. Here, we examined the propensities of three drug-like compounds, previously shown to bind and inhibit the cellular activity of three distinct RNAs, to non-specifically bind two HIV-1 stem-loop RNAs: the transactivation response element (TAR) and stem IIB in the rev response element (RREIIB). All three compounds bound to TAR and RREIIB *in vitro*, and two inhibited TAR-dependent transactivation and RRE-dependent viral export in cell-based assays while also exhibiting substantial off-target interactions consistent with non-specific cellular activity. A survey of X-ray and NMR structures of RNA-small molecule complexes revealed that drug-like molecules form hydrogen bonds with functional groups commonly accessible in canonical stem-loop RNA motifs, much like aminoglycosides, and in contrast to ligands that specifically bind riboswitches. Our results support extending the group of non-selective RNA-binders beyond aminoglycosides and intercalators to encompass drug-like compounds with capacity for non-specific hydrogen-bonding and reinforce the importance of assaying for off-target interactions and RNA selectivity *in vitro* and in cells when assessing novel RNA-binders.

## INTRODUCTION

RNA is an emerging class of attractive drug targets for a wide array of human diseases and pathogens (Connelly et al. 2016; Warner et al. 2018; Hermann 2016; Lieberman 2018; Matsui and Corey 2017). While there have been some successes in targeting RNAs with anti-sense oligonucleotides (ASOs) (Mendell et al. 2013; van Deutekom et al. 2013; Corey 2017), there is growing interest in developing small molecule inhibitors that can avoid delivery and safety limitations inherent to ASOs (Chi, Gatti, & Papoian, 2017; Gagnon & Corey, 2019; Geary, Norris, Yu, & Bennett, 2015; Orellana & Kasinski, 2017). Despite some success in identifying compounds that bind RNAs and inhibit their activities in cells and even in animal models (Costales et al. 2017; Ratni et al. 2018; Palacino et al. 2015; Parsons et al. 2009), many challenges remain in targeting RNA with small molecules. Chief among them is identifying small molecules that can bind to an intended RNA target while discriminating against all other cellular RNAs (Disney 2019; Thomas and Hergenrother 2008).

Achieving high selectivity when targeting RNA with small molecules is particularly challenging because unlike proteins with their twenty amino acids, RNAs are composed of only four chemically similar nucleotides, and their 3D structures are comprised of a smaller number of reoccurring motifs (Miao and Westhof 2017; Bevilacqua et al. 2016). Moreover, RNAs are highly susceptible to forming strong electrostatic and stacking interactions that are inherently non-specific. Indeed, aminoglycosides promiscuously bind RNA through electrostatic interactions (Wong et al. 1998; Walter et al. 1999; Verhelst et al. 2004) while intercalators bind to RNAs non-specifically through hydrophobic and π-stacking interactions (Tanner and Cech 1985; Tanious et al. 1992; White and Draper 1987). Both are known to have many side effects when used clinically due to non-specific RNA binding (Gunanathan Jayaraj et al. 2017; Hong et al. 2015; Xie et al. 2011; Callejo et al. 2015; Sharifi and Aragon-Ching 2012). Additionally, many RNA drug targets lack tertiary structure, and form highly flexible structures that can adaptively bind to a variety of small molecules (Bardaro et al. 2009; Hermann and Patel 2000; Stelzer et al. 2011).

Recent effort has focused on developing drug-like small molecules that minimize non-specific electrostatic and stacking interactions to target RNA (Hermann 2016; Warner et al. 2018; Childs-Disney and Disney 2016). These drug-like molecules are thought to primarily bind RNA through a combination of shape complementarity and hydrogen bonding (H-bonding) (Warner et al. 2018; Di Giorgio and Duca 2019). Such compounds with demonstrated cellular activity have been enumerated in the R-BIND database (Morgan et al. 2019). Some of these compounds have been shown to bind specific RNAs and inhibit their cellular activities with well-validated selectivity (Costales et al. 2017, 2019; Zhang et al. 2020; Shi et al. 2019; Haga et al. 2015; Naro et al. 2018). However, unlike aminoglycosides and intercalators, there are fewer in depth studies of selectivity for this class of drug-like RNA-binders particularly against stem-loop RNAs, which largely constitute the cellular transcriptome.

A common approach used to assess the *in vitro* RNA binding selectivity of drug-like compounds to stem-loop RNAs is to measure binding in the presence of excess tRNA or B-form DNA (Pascale et al. 2016; Ganser et al. 2018). However, neither tRNA nor B-DNA are good representatives of the structurally related and highly abundant RNA transcripts that compete for small molecule binding in the cell. Fewer studies have examined whether drug-like molecules can discriminate against simple stem-loop RNAs, which are more representative of the transcriptome (Ironmonger et al. 2007; Sztuba-Solinska et al. 2014; Velagapudi et al. 2014; Duca et al. 2010). When these selectivity tests are performed, some level of non-specific binding is often reported (Ironmonger et al. 2007; Duca et al. 2010; Sztuba-Solinska et al. 2014).

Similarly, in cell-based functional assays, controls to assess compound activity in the absence of the target RNA (off-target effects) or against a distinct RNA (cellular selectivity) are not always performed. When performed, drug-like RNA-targeted small molecules are often found to have broad, promiscuous activity (Schmidt 2014; Murchie et al. 2004; Mischiati et al. 2001; Nahar et al. 2014) and/or to interact with assay reporter proteins (Thorne et al. 2010). Because few studies comprehensively report on the selectivity of these drug-like molecules at both the *in vitro* and cellular level (Zhang et al. 2020; Mischiati et al. 2001; Richter et al. 2004), it is unclear what capacity this new class of RNA-targeted compounds has to bind RNAs non-specifically and what structural features define this behavior.

In this study, we evaluated the propensity of three compounds that are representatives of this new drug-like class of RNA binders to non-specifically bind stem-loop RNAs containing bulges and internal loops. The compounds were DPQ, pentamidine, and yohimbine (Fig. 1A), which were previously shown to bind and inhibit the cellular activity of three different RNAs: the influenza A virus (IAV) promoter (Lee et al. 2014), CUG repeats (Warf et al. 2009), and the ferritin iron-response element (IRE) (Tibodeau et al. 2006) respectively. We then assayed these small molecules for their activity against two unrelated RNAs that form stem-loop structures representative of the cellular transcriptome; the HIV-1 transactivation response element (TAR) (Puglisi et al. 1992) and stem IIB in the rev response element (RREIIB) (Malim et al. 1990, 1989; Chang and Sharp 1989; Malim et al. 1988) (Fig. 1B).

**Figure 1.**
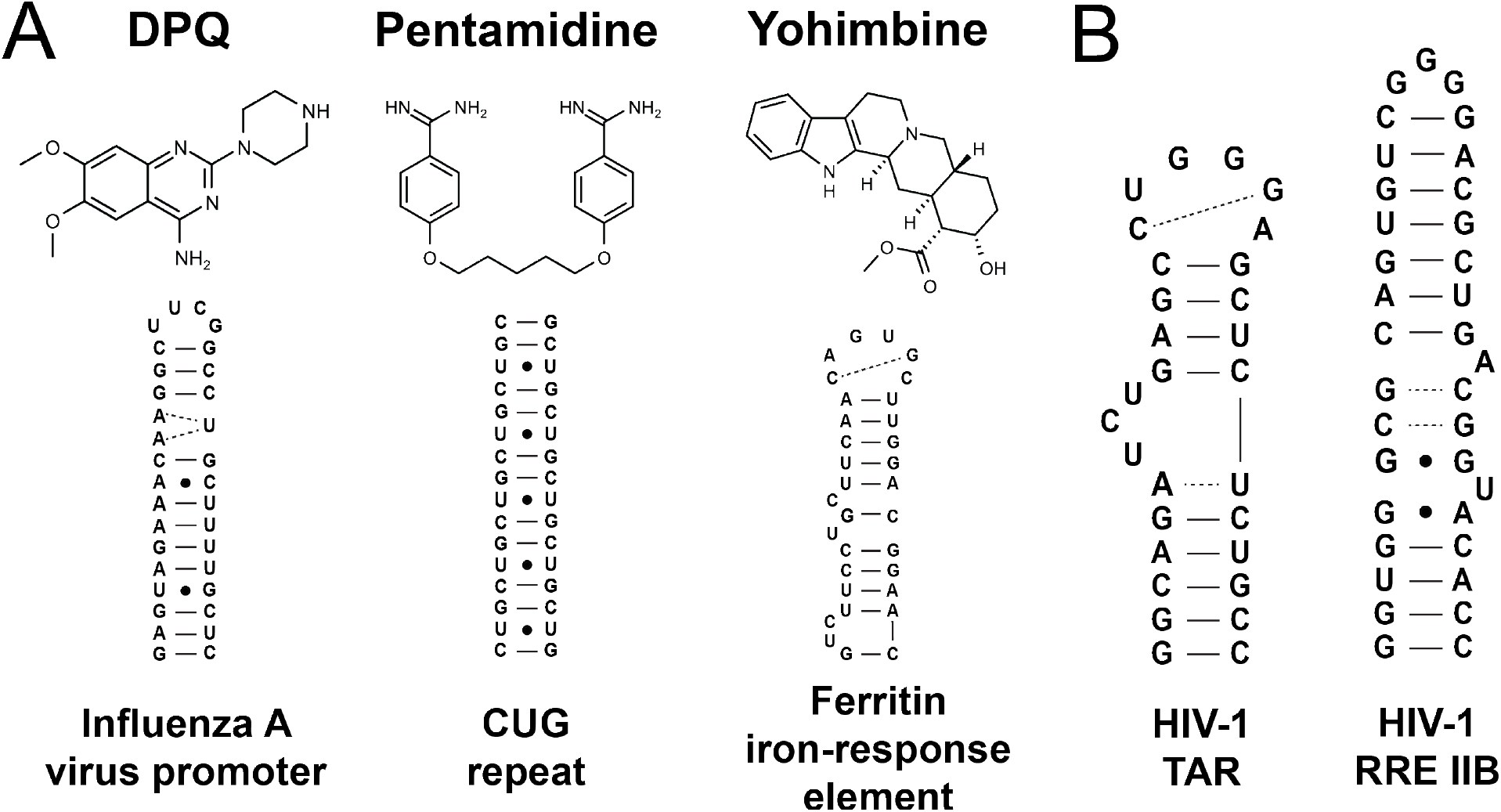
Small molecules and RNA constructs used in this study. *(A)* Structures of three RNA-binding small molecules from previous studies (Lee et al. 2014; Warf et al. 2009; Tibodeau et al. 2006) with the secondary structures of the RNAs they were previously reported to bind. *(B)* Secondary structures of HIV-1 TAR and RRE.

DPQ was previously shown (Lee et al. 2014) to bind the IAV promoter *in vitro* and to inhibit replication of IAV H1N1, IAV H3N2, and influenza B virus in cells with EC_50_ ~ 72-276 μM. Little is known about the RNA binding specificity or off-target effects of DPQ. Pentamidine, which is an FDA-approved drug for several antimicrobial indications, was shown to bind CUG RNA repeats and to inhibit MBNL1 binding *in vitro* with IC_50_ ~ 59 μM (Warf et al. 2009), and to actively rescue splicing of pre-mRNAs regulated by MBNL1 in a CUG-dependent manner in cell-based assays and an animal model. Interestingly, in the absence of CUG repeat RNAs, pentamidine was shown to have opposite effects and this was attributed to non-specific binding to an intron stem-loop (Warf et al. 2009). Finally, yohimbine was shown to weakly bind the ferritin IRE *in vitro* with K_d_ ~3.9 mM. Yet despite this weak binding affinity, yohimbine was shown to increase the rate of ferritin translation in a cell-free expression system and to discriminate against a ferritin IRE point mutant that has reduced affinity for an essential protein binding partner, indicating some selectivity for the native ferritin IRE RNA (Tibodeau et al. 2006).

Surprisingly, all three compounds bound to both TAR and RREIIB *in vitro*. For DPQ and pentamidine, the *in vitro* binding affinities were comparable to those reported for the original RNA targets (Table 1), and both compounds showed dose-dependent inhibition of TAR-dependent transactivation as well as RRE-dependent viral export in cell-based assays (Fig. 3). However, the compounds also showed clear off-target interactions in both cell-based assays indicating non-specific binding. NMR chemical shift mapping combined with a structure-based survey reveals that the drug-like small molecules form H-bonds with functional groups that are commonly accessible in stem-loop RNAs (Fig. 4). Our results show that even near neutral and non-planar drug-like compounds can promiscuously bind RNAs, most likely by H-bonding, and that non-specific RNA-binders can appear to specifically inhibit their cellular activities if measurements of off-target interactions and cellular specificity are not performed.

**Table 1.** IC_50_s and apparent K_d_s describing binding of each small molecule to TAR and RREIIB obtained from peptide displacement assays (see Methods). Uncertainty reflects the standard deviation over three independent measurements. When there was no sufficient change in signal to fit a binding curve, the binding constant was not determined (n.d.). Also shown is the K_d_ to other target RNAs reported previously in the literature (Lee et al. 2014; Warf et al. 2009; Tibodeau et al. 2006).

| Small molecule | IC<sub>50</sub> TAR (μM) | K<sub>d,app</sub> TAR (μM) | K<sub>d, app</sub> RRE-Rev complex (μM) | Binding affinity to target (μM) | Intended target |
| --- | --- | --- | --- | --- | --- |
| DPQ | 51 ± 4 | 43 ± 4 | 140 ± 55 | 51 ± 9 | IAV promoter |
| Pentamidine | 397 ± 82 | 332 ± 60 | >1000 | 58 ± 5 | CUG repeat |
| Yohimbine | >1000 | >1000 | n.d. | 3,900 ± 1,200 | Ferritin IRE |

## RESULTS

### Small molecules bind to TAR and RREIIB*in vitro*

We used solution state NMR spectroscopy to test binding of the three small molecules to TAR and RREIIB. 2D [^13^C, ^1^H] SOFAST-HMQC (Sathyamoorthy et al. 2014) experiments were recorded on uniformly ^13^C/^15^N labeled TAR or RREIIB following addition of the small molecule to RNA. Surprisingly, all three small molecules resulted in distinct chemical shift perturbations (CSPs) for resonances across TAR (Fig. 2A) and RREIIB (Fig. 2B), with and without Mg^2+^, consistent with binding (Fig. 2; Supplemental Fig. S1-S3). For all molecules, the CSPs were larger for TAR (Fig. 2A) compared to RREIIB (Fig. 2B), which may reflect the higher flexibility of TAR, and a greater propensity to adapt its conformation to optimally bind different small molecules (Stelzer et al. 2011; Bardaro et al. 2009; Pitt et al. 2005).

**Figure 2.**
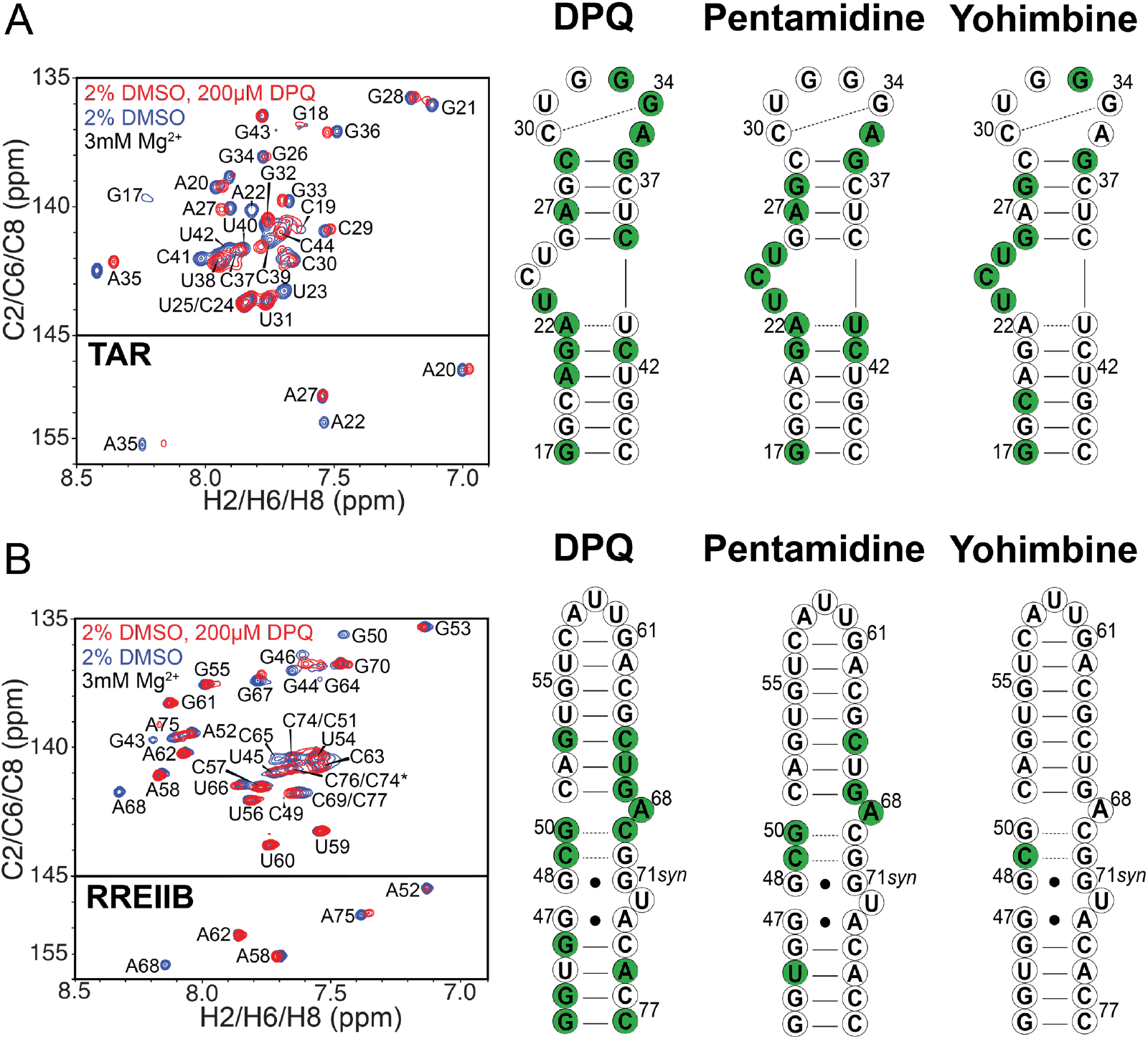
Testing small molecule binding to TAR and RREII using NMR chemical shift mapping experiments. Shown on the left are representative overlays of aromatic 2D [^13^C, ^1^H] SOFAST-HMQC (Sathyamoorthy et al. 2014) spectra for free and DPQ bound *(A)* TAR and *(B)* RREIIB showing chemical shift perturbations (CSPs) induced by DPQ. Buffer conditions were 15 mM NaH_2_PO_4_/Na_2_HPO_4_, 25 mM NaCl, 0.1 mM EDTA, 10% (v/v) D_2_O at pH 6.4 and 3 mM Mg^2+^ added directly to sample. Also shown on the right are the secondary structures of TAR and RREIIB in which residues that have <50% overlap between free and small molecule-bound spectra, or are absent in the bound spectra, are colored green.

**Figure 3.**
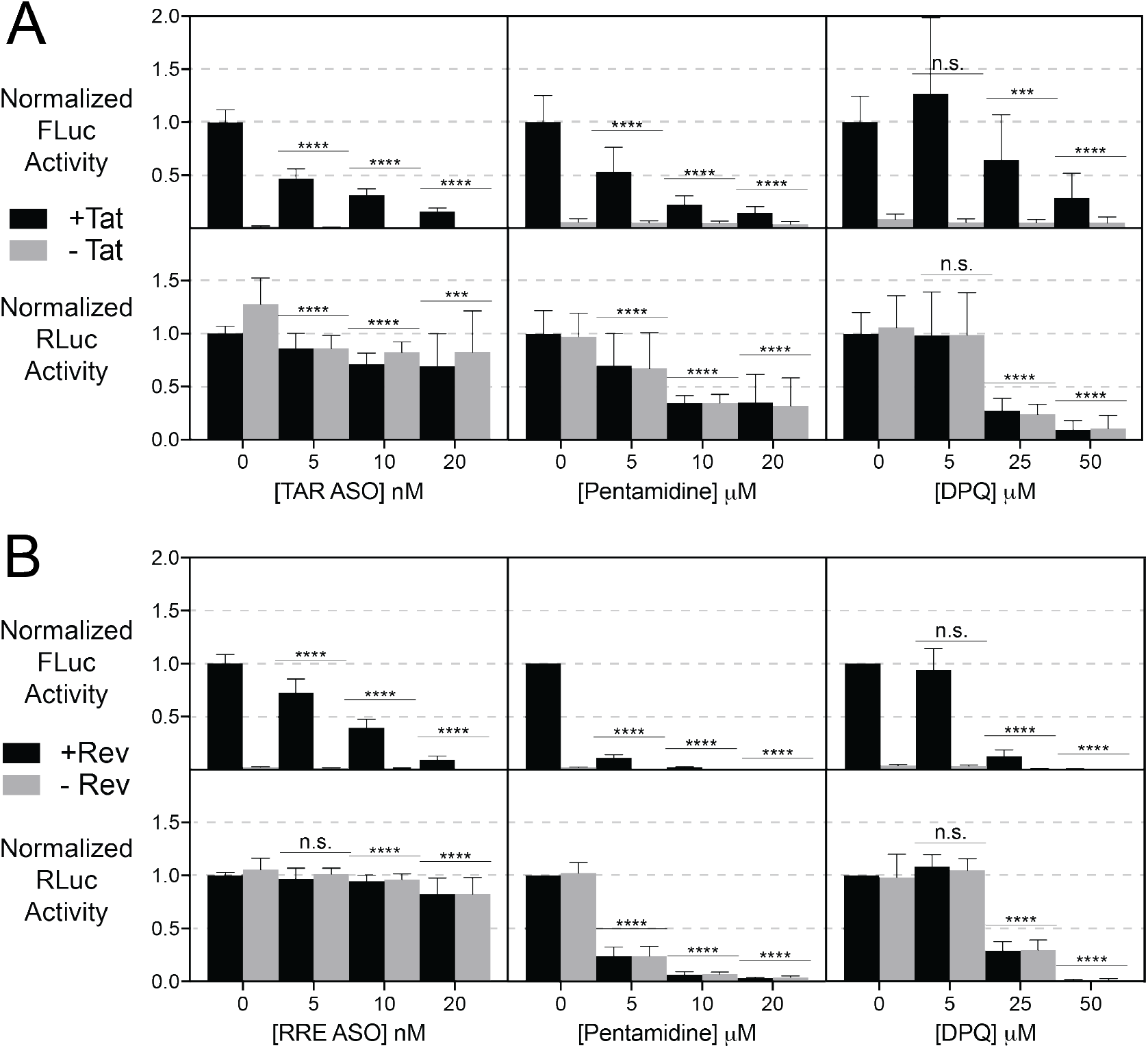
Cell-based functional assays for TAR and RRE in the presence of RNA-targeted ASO, Pentamidine, and DPQ. *(A)* Results for the Tat-dependent trans-activation assay. Top panels show FLuc activity, which is dependent on the TAR-Tat interaction. Bottom panels show RLuc activity, which is driven by a CMV promoter and is independent of the TAR-Tat interaction. Black bars indicate the activity in the presence of Tat, grey bars indicate activity in the absence of Tat. *(B)* Results for the Rev-dependent viral export assay. Top panels show FLuc activity, which is dependent on the RRE-Rev interaction. Bottom panels show RLuc activity, which is driven by a CMV promoter and is independent of the RRE-Rev interaction. Black bars indicate activity in the presence of Rev, grey bars indicate activity in the absence of Rev. n>5, with at least three biological replicates. *p<0.05 **p<0.01 ***p<0.001 ****p<0.0001 n.s. = no significance.

**Figure 4.**
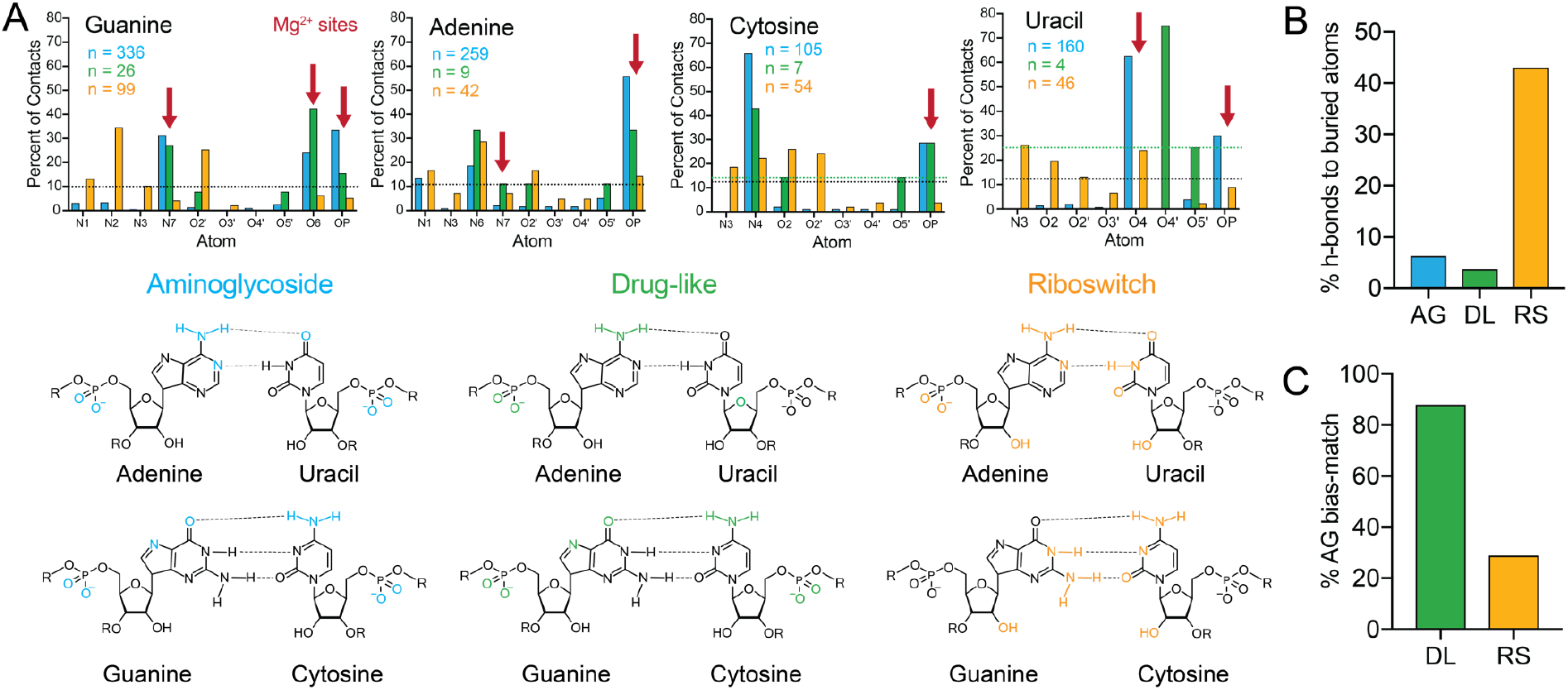
Structure-based survey of H-bonding in crystal and NMR structures of RNA-small molecule complexes. The data for aminoglycoside-RNA complexes (AG) are in blue, drug-like small molecule-RNA complexes (DL) in green, and small molecule-riboswitch complexes (RS) in orange. *(A)* Top: Column graphs representing the percentage of total H-bonds (n) to each nucleotide (A, U, G, and C) attributed to different atoms in the nucleotide. RNA atoms that are preferred Mg^2+^ binding sites (Zheng et al. 2015) are marked by red arrows. The black dotted line represents expected percentage of H-bond contacts if there is no bias (1 / # available H-bond donor and H-bond acceptor atoms for each unique nucleotide). Because of the small H-bond sample size (n=7 for cytosine, n=4 for uracil) for the drug-like subset, a green dotted line is also shown for uracil and cytosine representing the expected percentage of H-bond contacts if there is no bias for the drug-like group (1/7 for cytosine, 1/4 for uracil). Bottom: Chemical structure of Watson-Crick base-pairs in which atoms are colored if they exceed the bias threshold. *(B)* Percentage of H-bond contacts to atoms that are buried in a canonical RNA helix (AN1, AN3, UO2, UO3, GN1, GN2, GN3, CO2, CN3) for each group. *(C)* Percentage of contacts that pass the bias threshold for which the aminoglycoside group also passes the threshold. This is a measure of the similarity of binding patterns between the drug-like and riboswitch groups to the aminoglycoside group.

The CSPs were observed throughout the TAR and RREIIB molecules. While these could reflect conformational changes arising from small molecule binding at a distal site, a recent study employing TAR fragments showed that such CSPs arise from non-specific binding across the entire RNA molecule (Orlovsky et al. 2020). Interestingly, the CSPs were smaller and more localized in the presence of Mg^2+^, possibly because Mg^2+^ competes with the small molecules for H-bonding sites on the RNAs (Supplemental Fig. S1-S3) (Holbrook et al. 1977; Hermann and Westhof 1998). These data indicate that even near neutral non-planar compounds representative of drug-like class of RNA binders, bind to RNAs non-specifically.

### Assaying binding using RNA-peptide displacement assays

Next, we tested the ability of the compounds to displace peptides mimics of the cognate proteins that bind TAR and RREIIB using fluorescence-based assays. These assays have previously been used in screening for an evaluating TAR and RRE binders (Matsumoto et al. 2000; Hamasaki and Ueno 2001; Patwardhan et al. 2017, 2019; Zeiger et al. 2014). The TAR assay measures fluorescence resonance energy transfer (FRET) of a dual-labeled arginine rich motif (ARM) Tat-mimic peptide (Matsumoto et al. 2000; Ganser et al. 2018), and the RRE assay measures fluorescence anisotropy of an ARM Rev-mimic peptide (Luedtke and Tor 2003; Chu et al. 2019). We performed a dose-response displacement assay for all three compounds in the presence of TAR and Tat-ARM peptide and found that all have measurable IC_50_s, indicating binding to TAR and displacement of the Tat-ARM peptide (Table 1; Supplemental Fig. S4A). Interestingly, in the RRE Rev-ARM peptide displacement assay, DPQ and pentamidine increased rather than decreased the fluorescence anisotropy in a dose-dependent manner (Supplemental Fig. S4B). This indicates that the compounds bind to the Rev-RREIIB complex without displacing the Rev peptide, and possibly increase fluorescence anisotropy by changing the shape of the complex and/or aggregation. Indeed, the compounds did not affect the fluorescence anisotropy of the Rev-ARM peptide in the absence of RRE, which suggests that they are not causing aggregation of the peptide. Yohimbine showed no measurable change in fluorescence anisotropy (Table 1; Supplemental Fig. S4B), which is consistent with the very minor CSPs seen by NMR (Fig. 2B, Supplementary Fig. S1).

We calculated (see Methods) apparent K_i_s for all three compounds binding to TAR using the measured IC_50_s for RNA-small molecule binding and the K_d_s for RNA-peptide binding, (Table 1, Supplemental Fig. S5). For DPQ, the apparent K_i_ ~ 42 μM for TAR binding was comparable to the K_d_ ~ 51 μM reported for binding to its intended target RNA, the IAV promoter (Lee et al. 2014), while the K_i_ ~ 140 μM for binding RREIIB was 3-fold higher. Although different approaches were used to measure the binding affinities and the assays used different conditions, the binding affinity of DPQ for TAR and RREIIB clearly do not differ substantially from its target influenza A promoter.

For pentamidine, the estimated apparent K_i_ ~ 397 μM for TAR binding was 6-fold weaker than the IC_50_ ~ 58 μM reported for binding to CUG repeats (Warf et al. 2009), and for RREIIB it could not be reliably determined but we can estimate that K_d,app_ >1,000 μM. This suggests some degree of specificity for pentamidine binding to CUG repeats versus TAR and even more so RREIIB. Yohimbine’s K_i_ to TAR could not be reliably determined either, but we can estimate that it is also >1,000 μM, noting that K_d_ = 3,900 μM for binding to ferritin IRE (Tibodeau et al. 2006). Taken together, these results indicate that these drug-like small molecules can bind to unrelated stem-loop RNAs with comparable affinities.

### DPQ and pentamidine inhibit the biological activity of TAR and RRE in cell-based assays

We examined if the small molecules also inhibited the activity of TAR and RRE in cell-based assays (Ganser et al. 2020; Cullen 1986). To assess the effect of the small molecules on the activity of TAR in the cellular context, we transiently transfected HeLa cells with pFLuc-TAR (Ganser et al. 2020) in which TAR drives firefly-luciferase (FLuc) expression, and pcTat (Tiley et al. 1992) and pRLuc (Ganser et al. 2020), which are both driven by the constitutively active cytomegalovirus (CMV) immediate early promoter. When Tat is expressed it binds TAR and recruits host machinery to form the super elongation complex (SEC), allowing for transactivation of TAR and expression of FLuc. Expression of renilla-luciferase (RLuc) is not Tat-dependent and can be used to assay off-target activity. FLuc expression was also measured in the absence of Tat for every condition as a control for basal transcription. A small molecule specific for TAR is expected to show a decrease in FLuc activity in the presence of Tat, while having little to no effect on RLuc or the level of basal transcription in the absence of Tat.

As a positive control, we treated cells with a locked nucleic acid (LNA) antisense oligonucleotide (ASO) targeted to the bulge and loop region of HIV-1 TAR (Supplemental Table S1). We observed the expected dose-dependent decrease in FLuc activity with 50% expression at ~5 nM. The effect on RLuc was significantly smaller at all concentrations of the ASO and did not appear to be dose-dependent (Fig. 3A). Previous studies have also shown this type of marginal effect on an independent reporter with a similar sequence TAR-ASO (Turner et al. 2005). Next we tested the three compounds and found that two of them, DPQ and pentamidine, inhibited transactivation (50% decrease in FLuc activity in the presence of Tat at ~20 μM and ~5 μM respectively). However, both compounds also had a large, statistically significant dose-dependent effect on RLuc expression, consistent with off-target interactions possibly involving the inhibition of either transcription and/or translation in a TAR independent manner (Fig. 3A). Yohimbine did not show any effect on FLuc or RLuc in this assay, likely because of its low binding affinity for TAR (data not shown).

To de-convolute the contributions of TAR-dependent and TAR-independent drug interactions to the dose-dependent decrease in Fluc observed in the presence of Tat, we also measured the level of FLuc expression in the absence of Tat for each drug dose. The expression of FLuc in the absence of Tat also decreased with increasing drug concentrations, however a 2-way ANOVA analysis indicates that the drug-dependent FLuc decrease in the absence of Tat does not fully explain the FLuc decrease in the presence of Tat (Supplementary Table S1). These results indicate that while much of the inhibitory activity of the small molecule can be attributed to off-target interactions, some inhibition due to interactions between the small molecule and TAR cannot be ruled out.

To assess the effect of these molecules on RRE activity in the cellular context, we co-transfected 293T cells with the pcRev plasmid (Malim et al. 1988), the pFLuc-RRE plasmid (Ganser et al. 2020), and pRLuc (Ganser et al. 2020). When Rev binds and assembles on RRE, FLuc can be expressed due to Rev-RRE-mediated intron-containing mRNA export. RLuc expression was again used to assay for off-target interactions, and FLuc expression in absence of Rev was measured as a control for basal transcription. We used an RRE-targeted 20-mer LNA ASO as a positive control and observed a dose-dependent effect on viral export with 50% inhibition in the presence of Rev at ~5-10 nM (Fig. 3B). Again, the ASO had a minor effect on RLuc expression with statistically significant changes only observed at the highest concentrations.

When testing the three experimental compounds in this assay, we found that pentamidine and DPQ had a dose-dependent effect on RRE-dependent viral export (50% inhibition observed at ~10-20 μM and ~1-5 μM respectively). However, once again, the compounds also substantially decreased RLuc activity, indicating off-target interactions (Fig. 3B). We also observed decreased FLuc with increasing drug in the absence of Rev, but 2-way ANOVA analysis indicates that the drug-dependent FLuc decrease in the absence of Rev does not fully explain the FLuc decrease in the presence of Rev (Supplementary Table S1). Again, these results indicate that while most of the inhibitory activity of the small molecule can be attributed to off-target interactions, some inhibition due to interactions between the small molecule and RREIIB cannot be ruled out. Taken together, these data reveal that DPQ and pentamidine exhibit low cellular selectivity due to their inhibitory activity against both TAR and RRE in cells, and their dose-dependent effect on RLuc in both assays demonstrates substantial off-target interactions.

### High propensity for false positives in cell-based functional assays

Given that drug-like small molecules have a high propensity to non-specifically inhibit activity in cell-based functional assays, we surveyed the literature to assess how often specificity controls are performed. We examined all published studies reporting small molecules that bind HIV-1 TAR (Supplementary Table S2) and found that four out of ten studies reporting cell-based functional assays did not measure off-target interactions (Gelus et al. 1999; He et al. 2005; Hwang et al. 2003; Mischiati et al. 2004). Of those six studies that did, 33% of total compounds tested were found to have significant off-target interactions (Murchie et al. 2004; Mischiati et al. 2001; Hamy et al. 1998). Only one out of the ten studies measured and observed cellular selectivity by performing a cell-based functional assay for an RNA mutant (Stelzer et al. 2011).

One of the drug-like small molecules reported to inhibit TAR-dependent trans-activation in a cell-based assay, furamidine, was commercially available. In the original publication (Gelus et al. 1999) this compound was shown to inhibit transactivation in cells with IC_50_ ~ 30 μM, however measurements of off-target interactions were not performed. Additionally, furamidine had been shown in a previous study to bind RREIIB *in vitro* (Ratmeyer et al. 1996), indicating it may have nonspecific binding capacity and its activity in the transactivation assay may not be fully due to TAR-binding.

We tested furamidine in our cell-based Tat-dependent transactivation assay as described above. We found that furamidine, like pentamidine and DPQ, had a large dose dependent effect on both FLuc and RLuc (50% inhibition of FLuc ~20 μM), indicating that it also has abundant off-target effects that were not previously assessed (Supplemental Fig. S6). These results indicate that when the activity of the small molecule on the assay in the absence of the RNA of interest is not measured, many small molecules reported to inhibit RNA activity in cells may do so via off-target interactions that are unrelated to binding a target RNA.

### Drug-like molecules preferentially hydrogen bond to exposed sites in canonical stem-loop RNAs

Structural studies of DPQ bound to RNA (Lee et al. 2014) and of pentamidine bound to DNA (Edwards et al. 1992) show that they form H-bonds with functional groups in and around bulges, stems, internal loops, and mismatches. These include H-bonds between the methoxy oxygens of DPQ and the AUA internal loop of the IAV promotor, as well H-bonds with cytosine-N4 of the junctional G-C base pair (bp), and between the primary amine of pentamidine and adenine ribose-O4′. Because these are neutral, non-planar molecules and we observe H-bond contacts to sites commonly exposed and available for interaction in generic stem-loop RNAs composed of Watson-Crick bps, including TAR and RREIIB, we hypothesized that H-bonding is likely driving non-specific binding to RNAs by these drug-like molecules.

To test this hypothesis, we surveyed X-ray and NMR structures of stem-loop RNAs bound to drug-like small molecules (Dibrov et al. 2012; Davis et al. 2004; Lee et al. 2014) and enumerated all of the H-bonds between the small molecule and the RNA. As a negative control, we also surveyed the crystal structures of riboswitches (McCown et al. 2017; Schwalbe et al. 2007), which contain higher-order structural motifs and have evolved to bind metabolites with high selectivity. As a positive control, we surveyed the structures of RNAs bound to aminoglycosides (Faber et al. 2000; François et al. 2005; Kondo et al. 2007; Freisz et al. 2008; Han et al. 2005), which are well known to bind RNAs non-specifically (Fig. 4; Supplemental Table S3). It is interesting to note that there were not nearly as many drug-like small molecule-bound RNA structures (n=17) available as there were riboswitch (n=27) and aminoglycoside-RNA structures (n=60).

For aminoglycosides, ~94% of the H-bonds are with donor and acceptor atoms that are solvent exposed and accessible even in Watson-Crick bps (Fig. 4A-B), such as atoms in the phosphate backbone (OP, O5′), ribose moiety (O2′, O3′, O4′), and Hoogsteen face (N7, N4, O4) of the nucleobase (Walter et al. 1999). Guanine-N7, guanine-O6, uracil-O4, cytosine-N4, adenine-N6, and the phosphate oxygens (OP) for all bases are the positions most frequently involved in H-bonding (Fig. 4A). This pattern overlaps considerably with the preferred sites for Mg^2+^ association (Zheng et al. 2015; Ennifar et al. 1999), which include OP for all bases, guanine-N7, adenine-N7, uracil-O4, and guanine-O6 (Fig. 4A).

Interestingly, in stark contrast, the H-bonds present in riboswitch-ligand complexes are biased towards atoms residing in the Watson-Crick base-pairing face of the nucleobase. Approximately 43% of H-bonds were to RNA atoms that would otherwise be buried and inaccessible in a canonical RNA helix, such as guanine-N1, guanine-N2, cytosine-N1, cytosine-O2, and uracil-N3 (Fig. 4A,B). While we do not know whether metabolites can also form H-bonds with exposed atoms in Watson-Crick bps, nature has clearly evolved more complex binding pockets, which enable more specific ligand recognition (McCown et al. 2017; Schwalbe et al. 2007). One exception is the ribose 2′-OH group, which is highly represented riboswitch H-bonds.

Interestingly, we find that the drug-like small molecules, including DPQ, predominately model the behavior of aminoglycosides (Fig. 4C). 96% of H-bond contacts in the group of drug-like small molecules are formed with atoms that are solvent accessible in the canonical Watson-Crick bp helical structure (Fig. 4A,B), including guanine-N7 and the phosphate backbone (OP1 and OP2). Moreover, approximately 87% of the RNA sites of H-bonding overlap with those seen with aminoglycosides (Fig. 4C). This is in contrast with only 28% of riboswitch H-bonding sites overlapping with those observed with aminoglycosides (Fig. 4C). However, a notable departure from this behavior includes a slight bias towards H-bonding with ribose oxygens (O2′, O4′, O5′), similar to the riboswitch structures (Fig. 4A). This slight shift from binding to ribose oxygens versus phosphate oxygens may be due to the fact that these drug-like molecules have a near neutral charge, unlike aminoglycosides in which the positive charge may bias hydrogen bonding to the phosphate backbone. Therefore, the ability to form H-bonds with functional groups commonly presented in stem-loop RNAs provides a plausible mechanism for non-specific binding of drug-like small molecules to stem-loop RNAs.

## DISCUSSION

In concordance with observations previous studies (Schmidt 2014; Murchie et al. 2004; Mischiati et al. 2001; Nahar et al. 2014), our results extend the group of non-selective RNA binders beyond aminoglycosides and intercalators to encompass near-neutral, non-planar, drug-like compounds. These molecules likely bind RNAs non-specifically by H-bonding to exposed functional groups in and around canonical RNA motifs. The likelihood for non-specific binding is expected to be particularly high for low molecular weight compounds when targeting structurally simple stem-loop RNAs with few if any unique structural motifs, because they can only form a limited number of H-bonds that provide little discrimination against different but structurally related RNAs. Riboswitch ligands are also low molecular weight, but the RNA provides many unique binding motifs and geometries that increase the likelihood of specific binding. Indeed, work targeting CUG repeats and microRNAs (Rzuczek et al. 2017; Velagapudi et al. 2014) has shown that increased specificity towards stem-loop RNAs can be achieved with higher molecular weight compounds that increase the number of H-bonds. Thus, by increasing the H-bonding binding footprint and tailoring it to a target RNA, it is feasible to increase binding specificity to stem-loop RNAs.

Experiments that assess off-target interactions and RNA selectivity are clearly important when it comes to testing novel RNA-binders *in vitro* and *in vivo* (Naro et al. 2018; Zhang et al. 2020; Costales et al. 2017; Abulwerdi et al. 2019; Donlic et al. 2018). However, our survey of the HIV-1 TAR literature (Supplementary Table S2) revealed that these controls are not always performed (Gelus et al. 1999; He et al. 2005; Hwang et al. 2003). In fact, based on our survey of HIV-1 TAR studies, only six out of ten studies measured off-target interactions when reporting compounds with cell-activity, and those that did found 33% of the compounds to have promiscuous binding (Murchie et al. 2004; Mischiati et al. 2001; Hamy et al. 1998). Beyond TAR, studies of microRNA-targeted small molecules have shown that when off-target effects are assessed, many molecules have broad, promiscuous activity (Schmidt 2014; Nahar et al. 2014) and/or interact with assay reporter proteins (Thorne et al. 2010). Given the high tendency of drug-like small molecules with favorable H-bonding capacity to bind a variety of RNAs and to even have apparent cellular activity, controls that test off-target interactions and cellular specificity are of paramount importance.

Finally, our results show that non-specific RNA binders could have some degree of clinically useful broad-spectrum activity against different RNAs that may be overexpressed in different disease states. Not only does pentamidine bind model CUG repeats with 6-fold higher affinity *in vitro* compared to both TAR and RREIIB, CUG repeats are endogenously overexpressed in the diseased state. This may explain its efficacy in a mouse model of the disease (Warf et al. 2009). DPQ clearly inhibits influenza replication and it remains possible that much of this inhibition is the result of binding to the influenza A virus promoter. Indeed, in our own studies, both pentamidine and DPQ showed some level of TAR and RREIIB inhibition in cell-based assays. Therefore, these drug-like small molecules could offer good starting points for optimizing selectivity toward specific target RNAs through optimization of hydrogen bonding and other interactions.

## MATERIALS AND METHODS

### RNA sample preparation

^13^C/^15^N labeled HIV1-TAR and RREIIB for NMR studies was prepared by *in vitro* transcription using a DNA template containing the T7 promoter (Integrated DNA Technologies). The DNA template was annealed at 50 μM DNA in the presence of 3 mM MgCl_2_ by heating to 95°C for 5 minutes and cooling on ice for 1 hour. The transcription reaction was carried out at 37°C for 12 hours with T7 polymerase (New England Biolabs) in the presence of ^13^C/^15^N labeled nucleotide triphosphates (Cambridge Isotope Laboratories, Inc.). Unlabeled HIV1-TAR and RREIIB for *in vitro* displacement assays was synthesized with the MerMade 6 DNA/RNA synthesizer (Bioautomation) using standard phosphoramidite chemistry and 2’-hydroxyl deprotection protocols. Both labeled and unlabeled samples were purified using the same methodology, using 20% (w/v) denaturing PAGE with 8M urea and 1X TBE. RNA was excised and then electroeluted (Whatman, GE Healthcare) in 1X TAE buffer. Eluted RNA was then concentrated and ethanol precipitated. RNA was then dissolved in water to a concentration of ~50 μM and annealed by heating at 95°C for five minutes and cooling on ice for 1 hour. For NMR experiments, ^13^C/^15^N labeled RNA was buffer exchanged using centrifugal concentration (3 kDa molecular weight cutoff, EMD Millipore) into NMR buffer (15 mM NaH_2_PO_4_/Na_2_HPO_4_, 25 mM NaCl, 0.1 mM EDTA, 10% (v/v) D_2_O at pH 6.4). For *in vitro* assays, unlabeled TAR RNA was diluted to 150 nM in Tris-HCl assay buffer (50 mM Tris-HCl, 50 mM KCl, 0.01% (v/v) Triton X-100 at pH 7.4), and unlabeled RRE RNA was diluted to 180 nM in reaction buffer (30 mM HEPES pH= 7.0, 100 mM KCl, 10 mM sodium phosphate, 10 mM ammonium acetate, 10 mM guanidinium chloride, 2 mM MgCl_2_, 20 mM NaCl, 0.5 mM EDTA, and 0.001% (v/v) Triton-X100)

### Small molecule and antisense oligonucleotide (ASO) preparation

Small molecules were ordered in powder format from MilliporeSigma: DPQ (6,7-dimethoxy-2-(1-piperazinyl)-4-quinazolinamine) #R733466; Pentamidine isethionate salt #P0547; Yohimbine #49768. Pentamidine and Yohimbine were dissolved in water to 20 mM stocks. DPQ was dissolved in DMSO to a 20 mM stock. Locked nucleic acid (LNA) ASOs were ordered from Qiagen and dissolved in water to 100 μM. The TAR (16mer) sequence was 5’ +C*+T*+C*C*Cm*A*G*G*C*T*C*A*G*+A*+T*+C 3’ while RRE (20mer) was 5’ +G*+G*+C*+C*+T*G*T*A*C*C*G*T*C*A*G*+C*+G*+T*+C*+A 3’, in which (+), (*), and (Cm) indicate LNA, phosphorothioate linkage, and cytosine base with a 2′O-methyl modification, respectively.

### NMR Experiments

All NMR experiments were performed at 25°C on a Bruker 600 MHz spectrometer equipped with triple resonance HCN cryogenic probes. ^13^C/^15^N labeled RNA was exchanged into NMR buffer (15 mM NaH_2_PO_4_/Na_2_HPO_4_, 25 mM NaCl, 0.1 mM EDTA, 10% (v/v) D_2_O at pH 6.4). For spectra in the presence of magnesium, Mg^2+^ was added directly to the sample to a final concentration of 3 mM. NMR spectra were processed with NMRPipe (Delaglio et al. 1995) and visualized with SPARKY (Goddard and Kneller 2006). All molecules were soluble in water except DPQ, which was dissolved in DMSO. NMR spectra for free and small molecule bound RNAs were recorded in 2% DMSO for the DPQ panels in Figure 3, and Supplemental Figures S1 and S2. NMR samples were prepared by mixing the RNA (50 μM) with small molecules (DPQ and pentamidine) at a 1:4 molar ratio. A molar ratio of 1:60 was used for yohimbine to observe the CSPs shown in Figure 2 and Supplementary Figures S1 and S2 consistent with its lower affinity to its target RNA (Tibodeau et al. 2006). This titration is shown in Supplemental Figure S3.

### Fluorescence-based TAR-Tat displacement assay

The fluorescence-based displacement assay used a peptide mimic of Tat containing an arginine rich motif (ARM), an N-terminal fluorescein label, and a C-terminal TAMRA label (N-AAARKKRRQRRR-C, Genscript), and MerMade-synthesized unlabeled HIV-1 TAR. The peptide is highly flexible when free in solution, allowing the two terminal fluorophores to interact and quench the fluorescent signal (Matsumoto et al. 2000). However, upon binding to TAR the peptide becomes structured and the two fluorophores are held apart, allowing fluorescence resonance energy transfer from fluorescein to TAMRA. Thus, when a small molecule displaces the peptide, the fluorescence signal decreases. For this assay we used a concentration of 50nM TAR and 20nM Tat-ARM peptide because this ratio gave the maximum fluorescence signal. TAR and Tat-ARM peptide were incubated with serial dilutions of the small molecules in a 384-well plate for 10 minutes. The assay buffer consisted of 50mM Tris-HCl, 50 mM KCl, 0.01% (v/v) Triton X-100 at pH 7.4. Fluorescence was then measured in triplicate using a CLARIOstar plate reader (BMG Labtech) with a 485 nm excitation wavelength and 590 nm emission wavelength. The fluorescence data were fit using a four parameter variable slope dose-response model using GraphPad Prism (Prism 2019) and Equation 1,

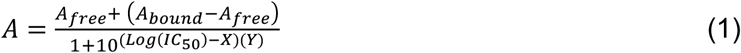

where A is the measured fluorescence at a given small molecule concentration (X); A_free_ is the measured fluorescence in the absence of TAR; A_bound_ is the fluorescence with saturated TAR-Tat binding; and Y is the Hill Slope. This assay was repeated three times, the average and standard deviation of the resulting 50% inhibitory constants (IC_50_) are reported in Table 1 and Supplementary Figure S4.

IC_50_ values were converted to apparent K_d_s using the Cheng-Prusoff Equation,

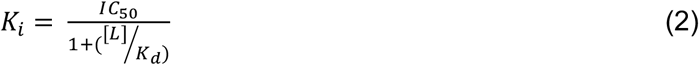

where K_i_ is the inhibition constant of the small molecule bound to TAR (Table 1); L is the constant concentration of Tat-ARM peptide used in determining the direct TAR-Tat K_d_ (20 nM); K_d_ is the binding constant between TAR and the Tat-ARM peptide (103.1 nM, Supplemental Fig. S5).

### Fluorescence-based RREIIB-Rev displacement assay

Fluorescence polarization displacement assays were carried out using 3’-end fluorescein labeled Rev-ARM peptide (Rev-Fl, TRQARRNRRRRWRERQRAAAACK-FITC, LifeTein LLC) (Chu et al. 2019). The serially diluted small molecule drugs in the reaction buffer (30 mM HEPES pH= 7.0, 100 mM KCl, 10 mM sodium phosphate, 10 mM ammonium acetate, 10 mM guanidinium chloride, 2 mM MgCl_2_, 20 mM NaCl, 0.5 mM EDTA, and 0.001% (v/v) Triton-X100) was incrementally added into a 384-well plate containing 10 nM Rev-Fl with or without 60 nM RREIIB (Prado et al. 2016; Chu et al. 2019). Fluorescence polarization (FP) was measured in triplicate using a CLARIOstar plate reader (BMG LABTECH) using 480 nm excitation and a 540 nm emission filter (Prado et al. 2016; Chu et al. 2019). This assay was repeated three times, the average and standard deviation of the resulting K_d,app_ values are reported in Table 1 and Supplementary Figure S4. These are K_d,app_ and not IC_50_ values because an increase in FP was observed, representing direct binding to the RREIIB-Rev peptide complex and not peptide displacement. The IC_50_ values were also fitted with the three parameter dose-response model in GraphPad Prism (Prism 2019) using Equation 3,

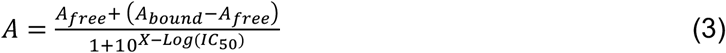

where A is the measured FP; A_free_ is the FP without Rev-Fl binding; A_bound_ is the FP with saturated Rev-Fl binding; X is the total small molecule concentration. K_d,app_ values were not converted to K_i_s because displacement of the peptide was not observed in the assay.

### TAR-Tat peptide binding assay

The fluorescence-based TAR-Tat peptide binding assay uses the same peptide and TAR construct as the displacement assay. In this assay a constant concentration of 20 nM Tat peptide is plated with serial dilutions of TAR in a 384-well plate. Both TAR and Tat peptide were diluted in assay buffer consisting of 50mM Tris-HCl, 100 mM NaCl, 0.01% (v/v) Triton X-100 at pH 7.4. Fluorescence was then measured in triplicate with a CLARIOstar plate reader (BMG Labtech) with a 485 nm excitation wavelength and 590 nm emission wavelength. This assay was repeated three times, the average and standard deviation of the K_d_ is reported in Supplemental Figure S6. Binding curves were fit to equation 4 in GraphPad Prism (Prism 2019) to determine the K_d_,

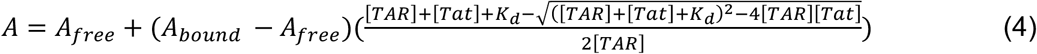

where A is the measured fluorescence; A_free_ is the fluorescence in the absence of TAR-Tat binding; A_bound_ is the fluorescence at saturated TAR-Tat binding; K_d_ is the measured apparent binding affinity and [TAR] and [Tat] are the concentrations of TAR and the peptide, respectively.

### TAR-Tat dependent trans-activation assay

pcTat (Tiley et al. 1992), pFLuc-TAR (Ganser et al. 2020), pcRLuc (Ganser et al. 2020), and pBC12-CMV (Tiley et al. 1992) expression plasmids were constructed as described previously. HeLa cells were maintained in Dulbecco’s modified Eagle medium (DMEM) supplemented with 10% fetal bovine serum and 0.1% gentamicin at 37°C and 5% CO_2_. Cells were plated to 1.5 × 10^5^ cells per well in 24-well plates, and treated with small molecule drug or vehicle only, 24 hours prior to transfection with polyethylenimine PEI (Polysciences). The primary transfection mixtures contained 250 ng pFLuc-TAR reporter plasmid, 10 ng RLuc control plasmid, + / − 20 ng pcTat expression plasmid, and pBC12-CMV filler DNA plasmid up to a total of 1510 ng total DNA per well. ASO-treated cells were transfected with ASO using PEI 10 minutes after transfection with the primary mixture. Media with drug or vehicle was replaced at 24 hours post-transfection, and cells were lysed at 4 hours post-transfection with 250 μL passive lysis buffer (Promega) and incubated 20 minutes at room temperature. FLuc and RLuc activity was measured using a Dual-Luciferase Reporter Assay System (Promega).

### RRE-Rev dependent export assay

pcRev (Malim et al. 1988), pFLuc-RRE (Ganser et al. 2020), pRLuc (Ganser et al. 2020), and pBC12-CMV (Tiley et al. 1992) expression plasmids were constructed as described previously. 293T cells were maintained in Dulbecco’s modified Eagle medium (DMEM) supplemented with 10% fetal bovine serum and 0.1% gentamicin at 37°C and 5% CO_2_. Cells were plated to 1 × 10^5^ cells per well in 24-well plates, and treated with small molecule drug or vehicle only, 24 hours prior to transfection with PEI. The primary transfection mixtures contained 5 ng pFLuc-RRE reporter plasmid, 5 ng RLuc control plasmid, 1 ng pcRev expression plasmid, and pBC12-CMV filler DNA plasmid up to a total of 1010 ng total DNA per well. ASO-treated cells were transfected with ASO using PEI 10 minutes after transfection with the primary mixture. Media with drug or vehicle was replaced at 24 hours post-transfection, and cells were lysed at 4 hours post-transfection with 250 μL passive lysis buffer (Promega) and incubated 20 minutes at room temperature. FLuc and RLuc activity was measured using a Dual-Luciferase Reporter Assay System (Promega).

### Statistical Analysis

Statistical analysis was done using the program JMP (JMP Pro, Version 14, SAS Institute Inc., Cary, NC, 1989-2019). For both the TAR-Tat dependent transactivation assay and the RRE-Rev dependent export assay, the raw luminescence values of each condition in each experiment were normalized to the average of the vehicle-treated, +Tat control luminescence values of that experiment. n = at least 6 replicates for each treatment group, with at least 3 being biological replicates. For each concentration of the small molecule or ASO, the + and - Tat conditions were compared to the + and -Tat conditions of the vehicle treated control in a 2-way ANOVA. The statistical significance of the main effect of the treatment on both the + and -Tat conditions are shown as asterisks above each condition in Figure 4. *p<0.05 **p<0.01 ***p<0.001 ****p<0.0001 n.s. = no significance. The p values of the interaction effect between Tat and drug are shown in supplemental table S1.

### TAR-binding small molecule literature survey and analysis

We surveyed studies that reported on small molecules binding to TAR published between 199 and 2020. From 1995-2014, we used the TAR-binding small molecule studies previously reported in Ganser et al, 2018. For the studies from 2014-2019, we performed a PubMed and Google Scholar search using the terms HIV AND TAR AND RNA AND binding with a 2014-2019 date filter. From the search results, we included all studies reported small molecules (not proteins or peptide mimetics) binding to TAR *in vitro* with a measurable IC_50_. We re-did the search replacing the word “binding” with “inhibit” and added any additional studies that fulfilled our criteria that were not included in the first search. There are 47 primary scientific articles included, however two of them we have classified into one study, as one article reported on *in vitro* and some cell-based experiments and a later article by the same authors included additional cell-based and viral studies of the same compounds (Mei et al. 1997, 1998). In total there are 46 studies, and they are enumerated in Supplemental Table 2. At the *in vitro* level we describe the assays used to measure affinity and any tests of RNA selectivity that were performed. At the cell based and viral assay levels we describe the experiments performed in each study as well as any tests of off-target interactions, RNA selectivity, cell viability, and list the cell lines and viral strains used.

### Structure Survey

This survey was conducted using the RCSB Protein Data Bank (PDB) in August of 2017 (Berman et al. 2007, 2000). We first constructed a master-database of all non-redundant RNA-ligand complexes (Supplemental Table S3). The PDB was filtered to only include structures that contain RNA, at least one ligand, and do not contain DNA or protein. A total of 623 X-ray and 58 NMR solution structures satisfying the above criteria were downloaded from the PDB website (https://www.rcsb.org). This set of structures was filtered to exclude structures in which the only type of small molecule ligand is an ion, a solvent molecule, or a linker molecule typically used to improve crystallization conditions. This reduced the dataset to 288 crystal structures and 48 NMR solution structures. This set of structures were filtered to exclude redundant structures that may bias the dataset. For any clusters of structures with global RMSDs < 2 Å (often the same RNA-drug pair), we included only one parent structure from that cluster. All RNA-ligand intermolecular H-bonds were identified for each structure using X3DNA-DSSR (Lu et al. 2015). We removed any PDB structures that did not include any hydrogen bonds (complex formed entirely by stacking interactions).

We then refined the H-bonding criteria for our master-database. X3DNA-DSSR has a very large range of distances between donor and acceptor atoms (4 Å and below), and so we applied additional distance criteria (between 2.0 Å and 3.5 Å) to more rigorously define H-bonds. To remove any additional sources of overrepresentation bias in the dataset, we removed all H-bonds that were redundant due to multiple identical bioassemblies within a single structure. All remaining H-bonds were then manually inspected to remove obvious false positive H-bonds such as donor-donor and acceptor-acceptor pairs, even considering potential tautomerizations. The resulting master-database includes a final set of 223 unique PBD structures (185 crystal and 38 NMR), comprised of 2,168 unique H-bonds in total (Supplemental Table S3).

We then refined this master-database to a specific subset for the purposes of this study, the data shown in Figure 4. First, we classified the PDB structures into three subgroups (1) stem-loop RNAs bound to aminoglycosides – 55 structures (2) riboswitches bound to their native ligand – 27 structures (3) stem-loop RNAs bound to drug-like small molecules – 17 structures for a total of 99 structures (71 crystal, 28 NMR) (Supplemental Table S4). Any PDB structures that did not fit into one of these three categories, such as synthetic aptamers with tertiary structure and complexes in which the small molecules are nucleotides that exhibit Watson-Crick base-pairing with the RNA to form a canonical helix, were excluded from this study. We then further refined the H-bonds in this subset of PDB structures. Any H-bonds to noncanonical (modified) RNA bases were excluded from analysis. Any palindromic RNA-small molecule interactions in a single structure were marked such that only a single copy of the hydrogen bonds was included in the analysis. These exclusions left a total of 1147 H-bonds from the 99 structures of this subset to be analyzed. The number of ligand H-bond contacts to each unique H-bond donor/acceptor RNA atom was then determined for each of the three PDB subgroups (aminoglycoside, drug-like, riboswitch) (Figure 4).

## ACKNOWLEDGEMENTS

We would like to thank the Duke Magnetic Resonance Spectroscopy Center for nuclear magnetic resonance resources. We would also like to thank Dr. Christopher Holley for helpful guidance on antisense oligonucleotide design and implementation, and Atul Rangadurai for critical reading of the manuscript. This work was supported by the National Institutes of Health (NIH) grants U54 AI150470 to H.M.A.-H. and F30 AI143282-01A1 to M.L.K.

