## Supplemental Materials for "Revealing the high propensity of RNAs to non-specifically bind drug-like small molecules"

### Supplementary Tables

| Assay | 5nM ASO | 10nM ASO | 20nM ASO | 5μM Pent | 10μM Pent | 20μM Pent | 5μM DPQ | 25μM DPQ | 50μM DPQ |
|---|---|---|---|---|---|---|---|---|---|
| TAR-Tat | <0.0001 | <0.0001 | <0.0001 | <0.0001 | <0.0001 | <0.0001 | 0.0545 | 0.0013 | <0.0001 |
| RRE-Rev | <0.0001 | <0.0001 | <0.0001 | <0.0001 | <0.0001 | <0.0001 | 0.5662 | <0.0001 | <0.0001 |

**Supplemental Table S1.** P values of the interaction between the drug condition and protein (Tat or Rev) condition for each drug dose. A 2-way ANOVA was performed to determine if there was a significant difference between the fold change of the +Tat and -Tat condition between the no drug control and each drug dose.

**Supplementary Table S2.** Literature survey of TAR-binding small molecules provided in attached excel sheet.

**Supplemental Table S3.** Complete structural survey of all non-redundant RNA-ligand complexes with all hydrogen bonding information, provided in attached excel sheet.

**Supplemental Table S4.** Subset of PDB structures from the structural survey used in this study, provided in attached excel sheet.

### Supplementary Figures

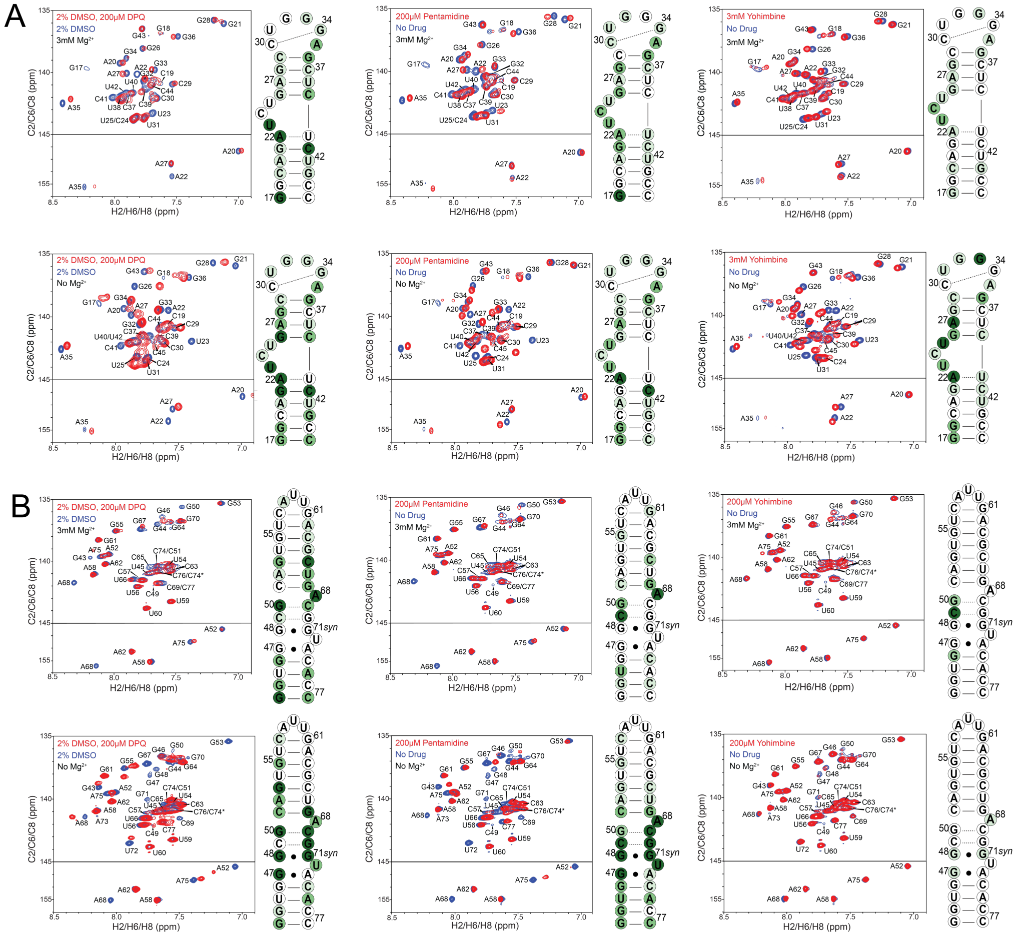

**Supplementary Figure S1.** The three small molecules bind to TAR and RREII based on NMR chemical shift mapping experiments. Shown are representative chemical shift perturbations between free and bound RNAs in aromatic 2D [$^{13}$C, $^{1}$H] SOFAST-HMQC (Sathyamoorthy et al. 2014) spectra for DPQ, pentamidine, and yohimbine (buffer: 15 mM $NaH_2PO_4/Na_2HPO_4$, 25 mM NaCl, 0.1 mM EDTA, 10% (v/v) $D_2O$ at pH 6.4; 3 mM $Mg^{2+}$ added directly to sample) for *(A)* TAR and *(B)* RREIIB in the absence and presence of 3 mM $Mg^{2+}$. Also shown to the right of each spectrum is the corresponding secondary structure of TAR or RREIIB in which residues that have <50% overlap between free and small molecule-bound spectra, or are absent in the bound spectra, are colored green.

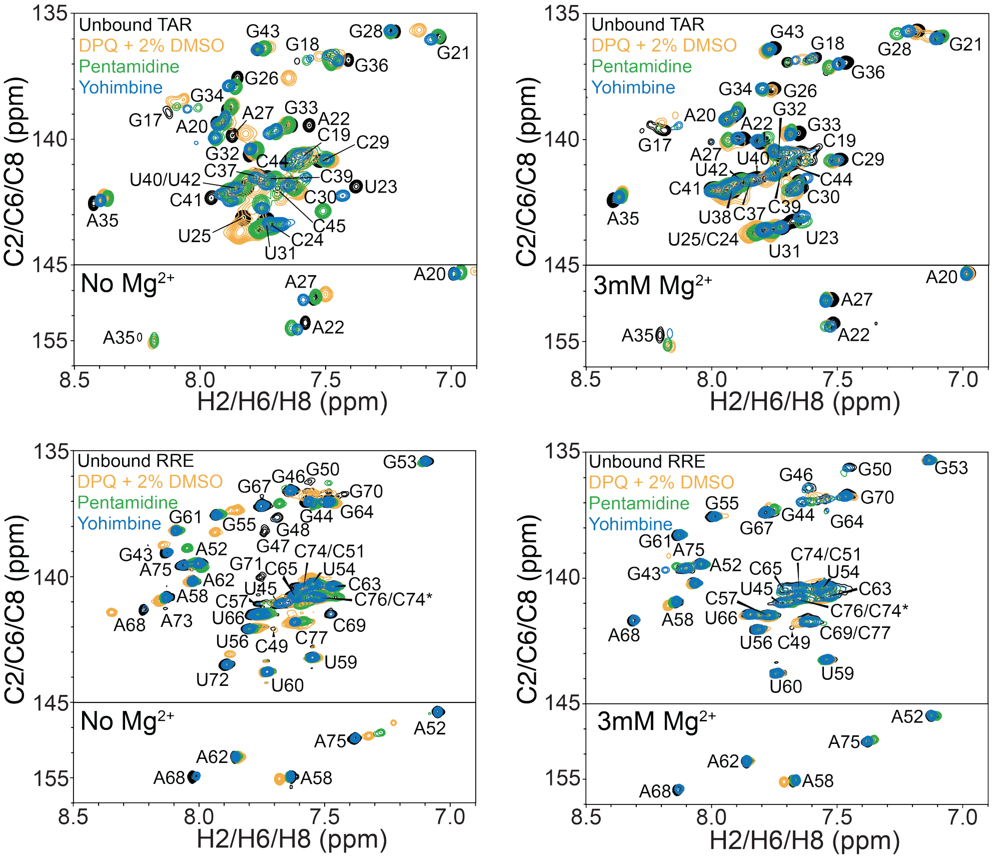


**Supplemental Figure S2.** Effect of $Mg^{2+}$ on 2D [$^{13}C$, $^{1}H$] SOFAST-HMQC (Sathyamoorthy et al. 2014) small molecule bound TAR and RREIIB. The magnitude of the CSPs are diminished for both TAR and RREIIB in the presence of $Mg^{2+}$.

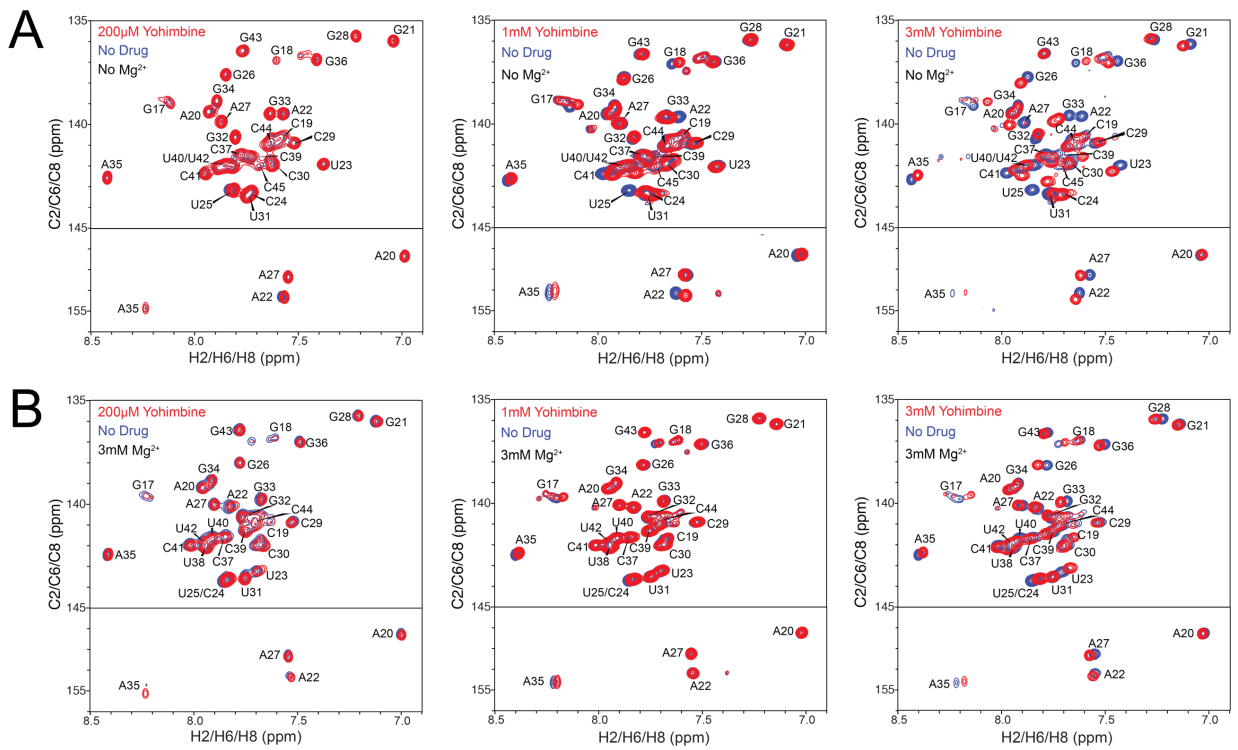


**Supplemental Figure S3.** 2D [$^{13}$C, $^{1}$H] SOFAST-HMQC (Sathyamoorthy et al. 2014) spectra of yohimbine titration with TAR in the absence *(A)* and presence *(B)* of 3 mM $Mg^{2+}$. The TAR and RREIIB concentrations were both 50 μM. Yohimbine was present at a 1:4 ratio (200 μM) in the left column, 1:30 ratio (1 mM) in the middle column, and 1:60 ratio (3 mM) in the right column.

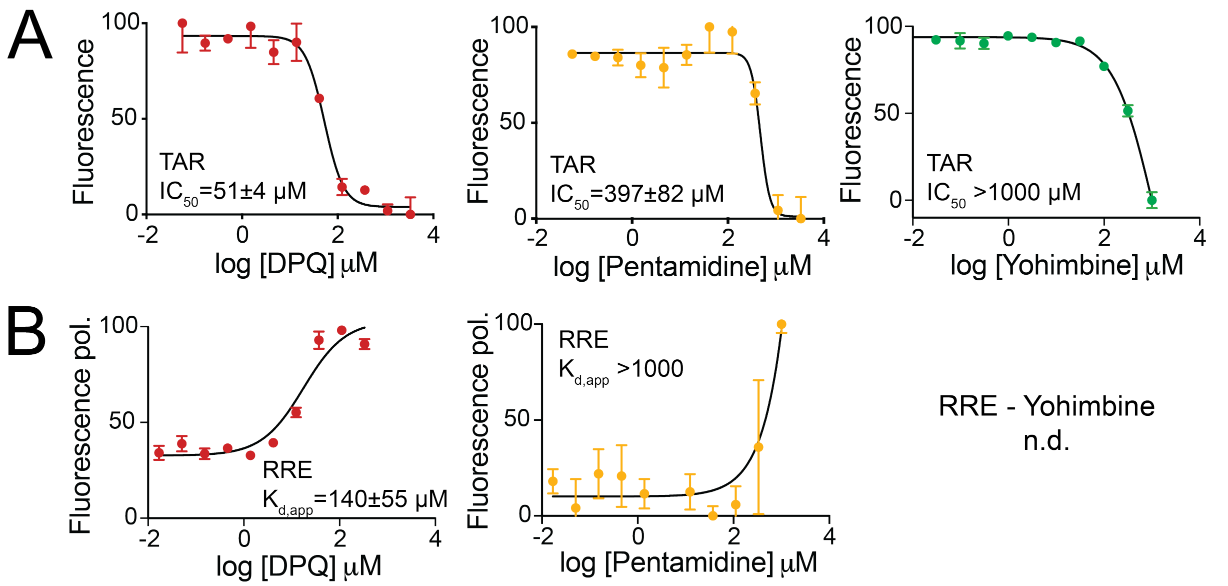


**Supplementary Figure S4.** Binding titrations used to measure $IC_{50}$s for small molecules binding to TAR and RRE. *(A)* FRET-based tat-peptide displacement assay. Shown is normalized fluorescence values measured for TAR following subtraction of baseline fluorescence in the absence of Tat-peptide, fitted with a variable-site binding model (See methods). Uncertainty reflects the standard deviation from three independent measurements *(B)* Shown is normalized fluorescence anisotropy values measured for TAR following subtraction of baseline fluorescence anisotropy in the absence of Rev-peptide, fitted with a one-site binding model (See methods). Uncertainty in the plotted data reflects the standard deviation from three independent measurements. The uncertainty in the $IC_{50}$s/$K_{d,app}$s reflects the standard deviation of three independent measurements.

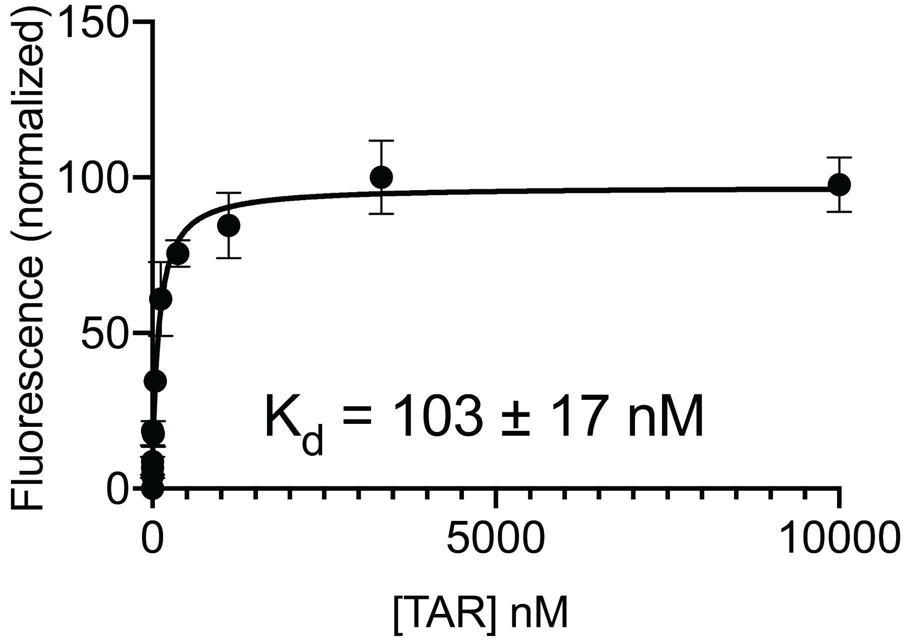


**Supplemental Figure S5.** Measurement of the binding affinity between Tat-peptide mimic and TAR using FRET-based fluorescence. Shown is normalized fluorescence values measured for TAR, fitted with a one-site binding model (See methods). Uncertainty in the plotted data reflects the standard deviation from three replicate measurements. The uncertainty in the $K_d$ value reflects the standard deviation of three independent measurements.

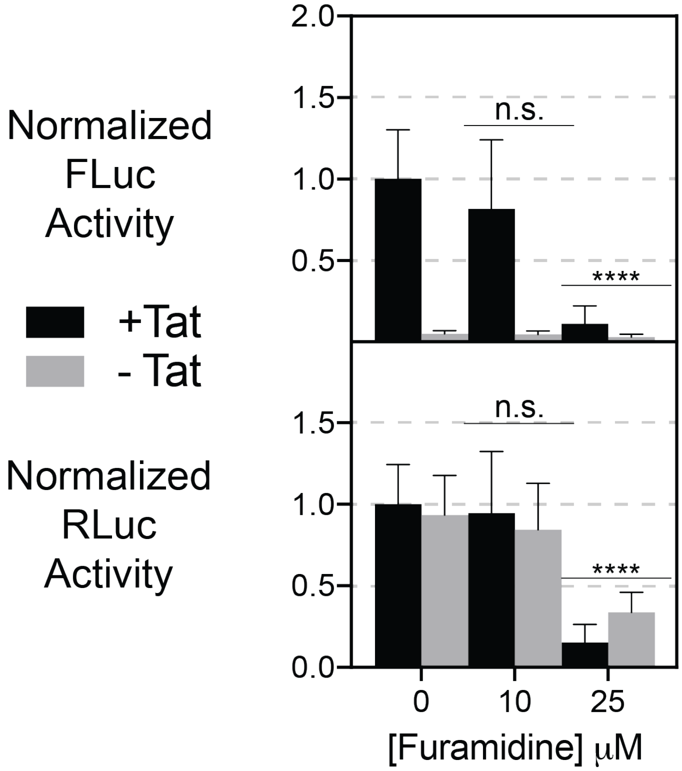


**Supplemental Figure S6.** Effect of furamidine on TAR-Tat dependent transactivation. Top panel indicates FLuc activity, which is dependent on the TAR-Tat interaction. Bottom panel indicates RLuc activity, which is driven by a CMV promoter and is independent of the TAR-Tat interaction. Black bars indicate activity the presence of Tat, grey bars indicate activity in the absence of Tat. n=at least 6, with at least 3 biological replicates. *$p<0.05$ **$p<0.01$ ***$p<0.001$ ****$p<0.0001$ n.s. = no significance.
